# Augmentation of nonsense mediated decay by rapamycin

**DOI:** 10.1101/028332

**Authors:** Rocio Teresa Martinez-Nunez, Doyle Coyne, Linnea Jansson, Miles Rush, Hanane Ennajdaoui, Tilman Sanchez-Elsner, Jeremy R. Sanford

**Affiliations:** University of California Santa Cruz, Department of Molecular, Cellular and Developmental Biology, Santa Cruz, CA 95064; Clinical and Experimental Sciences, Faculty of Medicine, University of Southampton, Southampton, SO16 6YD, UK

## Abstract

RNA surveillance by the Nonsense Mediated Decay (NMD) pathway eliminates potentially deleterious transcripts containing Premature Termination Codons (PTCs). The transition from a pioneering round of translation to steady state translation is hypothesized to be a major checkpoint in this process. One hallmark of mRNAs licensed for translation is the exchange of 7-methylguanosine cap binding proteins. However, mRNAs undergoing steady state translation are also NMD substrates, raising mechanistic questions about the NMD checkpoint. To test the role of cap binding proteins in NMD, we modulated the protein composition of cytoplasmic messenger ribonucleoprotein particles (mRNPs) with the naturally occurring macrolide rapamycin. We demonstrate that despite well-documented attenuation of cap-dependent mRNA translation, rapamycin can augment NMD. Rapamycin-treatment significantly reduces the levels of endogenous and exogenous PTC-containing mRNA isoforms in a dose- and UPF1- dependent manner. PTC-containing transcripts exhibit a shorter half-life upon rapamacyin-treatment as compared to non-PTC isoforms. Rapamycin also causes depletion of PTC-containing mRNA isoforms from polyribosomes, suggesting that actively translating ribosomes can transition between low and high NMD states. Importantly, mRNPs show depletion of eIF4E and retention of the nuclear Cap Binding Complex (CBC) in rapamycin-treated cells. Our data demonstrate that rapamycin potentiates pioneer-like mRNP context thereby decreasing NMD evasion.

## INTRODUCTION

Nonsense Mediated Decay (NMD) is a conserved RNA surveillance system which recognizes and eliminates transcripts containing Premature Termination Codons (PTCs) (1). The NMD system prevents the synthesis of potentially toxic proteins arising from transcripts containing nonsense mutations, frame shifts and those generated by alternative splicing. NMD is known to regulate the expression of many genes, including those involved in the NMD pathway itself (2,3).

The constellation of proteins associated with RNA transcripts is dynamic and reflects the diverse cellular landscapes traversed by the mRNA. For example, all newly synthesized mRNAs are bound by the nuclear Cap Binding Complex (CBC), the Exon Junction Complex (EJC) and the nuclear poly(A) binding protein (PABPN1) (4-6). By contrast, mRNAs licensed for translation are depleted of EJC proteins, and have exchanged CBC and PABPN1 at the cap and poly(A) tail for the eukaryotic translation initiation factor 4E (eIF4E) and the cytoplasmic poly(A) binding protein (PABPC), respectively (6-8). Termination codons located more than 25-35 nt upstream from an Exon Junction Complex (EJC) are strong NMD signals (9). Following PTC detection by a pioneering ribosome, the SMG1 phosphatidylinositol 3-kinase-related kinase (SMG1) and the EJC protein UP-Frameshift Mutation 1 homolog (UPF1) interact with the translation termination release factor-1 and -3 (eRF1 and eRF3, respectively) to form the SURF complex (10,11). The SURF complex associates with the downstream EJC and triggers SMG1-dependent phosphorylation of UPF1, which is required for inhibition of translation initiation and subsequent nucleolytic decay of the targeted messenger RNA (12). Additionally, shorter 3’UTRs are thought to suppress NMD and presence of *cis*-acting elements in long 3’UTRs can trigger NMD evasion (13,14).

NMD efficiency may be linked to the composition of messenger ribonucleoprotein particles (mRNPs). The CBC is displaced by eIF4E in a translation-independent manner (8,15). This exchange mechanism is driven by interactions of CBP80 with the nuclear import receptor Importin b (IMPb) and also presumably by mass action due to the high cytoplasmic concentrations of eIF4E (8,16). These rearrangements have important consequences for NMD; for example the interaction of CBC with UPF1 promotes assembly of the SURF complex and is important for activating NMD (17). Recent observations suggest that NMD in mammalian cells can also occur during eIF4E-initiated translation, in contrast to the pioneer round model (18,19). However these data raise the intriguing hypothesis that NMD efficiency and regulation might differ when triggered in the context of CBC-or eIF4E-bound mRNPs.

The mechanistic Target of Rapamycin Complex 1 (mTORC1) serves as part of a nutrient/stress sensing pathway and is inhibited by the naturally occurring macrolide rapamycin, which promotes autophagy and extends lifespan (20,21). One important branch of the mTORC1 signaling pathway modulates initiation of cap-dependent translation through phosphorylation of targets such as 4EBPs (eIF4E-Binding proteins) and S6K1 (40S ribosomal subunit protein S6 protein kinase 1). The 4EBP family functions as inhibitor of cap dependent translation by inducing nuclear sequestration of eIF4E (16). Active mTORC1 phosphorylates 4EBPs and counteracts their inhibitory effects on eIF4E (22) thereby stimulating translation. Interestingly, mTORC1 inhibition does not lead to global repression of translation but rather to a regulated inhibition of translation and control of gene expression (23). Furthermore, eIF4G, a translation initiation factor that is stimulated by mTOR, suppresses NMD (24). These observations prompted us to hypothesize that mTORC1 inhibition by rapamycin may enhance NMD.

Here we show that rapamycin augments decay of endogenous and exogenous PTC-containing mRNA isoforms in a dose and UPF1-dependent manner. Interestingly, PTC-containing mRNA isoforms accumulate in polyribosomes when mTORC1 is active, but are depleted when cells are treated with rapamycin. Because rapamycin inhibits mTORC1 causing nuclear sequestration of eIF4E, we hypothesized that rapamycin would alter the dynamics of CBC-and eIF4E-cap binding decreasing NMD evasion. Indeed, we observe that rapamycin treatment induces nuclear accumulation of eIF4E. Concomitantly, we show that rapamycin promotes eIF4E depletion from cytosolic mRNPs whilst binding of CBC is moderately increased, suggesting accumulation of pioneer-like mRNPs. Taken together, our results demonstrate that mammalian cells can dynamically regulate NMD efficiency of messenger RNA associated with polyribosomes depending on the cellular needs.

## MATERIAL AND METHODS

### Cell culture and Rapamycin treatment

Human Embryonic Kidney (HEK) 293 T cells were cultured in DMEM 10%FBS (Santa Cruz Biotech) at 37°C 5% CO_2_. For rapamycin (Cell Signaling) experiments, cells were starved for 24h in Glutamine deficient media DMEM 10%FBS. Following this, they were incubated in 22nM rapamycin or vehicle (DMSO) for 30min prior to feeding with complete medium containing DMSO or rapamycin. For polyribosome experiments, cells were harvested 8h-post rapamycin stimulation. For dose-response assays cells were harvested 72h at the indicated concentrations diluted in vehicle (DMSO). Actinomycin D was added after 48h rapamycin exposure at a concentration of 5mg/mL and cells harvested at the indicated time points.

### Immunofluorescence

HEK 293 T cells were seeded at low density on coverslips overnight. Cells were treated as described above, fixed with 3.2% paraformaldehyde for 15 min at room temperature and permeabilized with 0.1% Triton X-100 in PBS for 3 min. After blocking in 1% BSA+PBS 1h at room temperature they were incubated with eIF4E-FITC primary conjugated antibody (Santa Cruz Biotechnology) overnight at 4°C and washed. After washing 3 times with 0.02% tween PBS, coverslips were mounted using DAPI-mounting media.

### Microscopy and quantification

Immunofluorescence was observed using a Leica DM2700 white field microscope at magnification 63X (oil immersion objective). Quantification of fluorescence was performed using ImageJ.

### Cell fractionation and polyribosome extraction

Cells were incubated in the presence of 100mg/,mL cycloheximide 10min at 37C, 5%CO_2_ prior harvesting. Cell lysis was performed in 0.5% NP40, 20mM Tris HCl pH 7.5, 100mM KCl and 10mM MgCl_2_. Lysates were passed 3 times through a 23G needle and incubated on ice 7min. Extracts were then centrifuged at 10K rpm at 4C 7min. Cytosolic extracts were laid on 15-45% sucrose gradients prepared in 20mM Tris HCl pH 7.5, 100mM KCl and 10mM MgCl_2_ using a Gradient Station Master (Biocomp). Gradients were ultracentrifuged at 40K rpm 4C for 1h 20min using SW41 rotor in a Beckman Ultracentrifuge. Polyribosome monitoring and extraction of discrete complexes were conducted using the Gradient Station Master (Biocomp).

### Western blotting

Cell extracts were prepared in Lysis Buffer as described above. For 4E-BP and P-4E-BP blots, cell extracts were resolved in 12% Bis/acrylamide gels. For CBP80/eIF4E blots, extracts were resolved in 10% Bis/acrylamide gels. Antibodies used were: 4E-BP1 53H11 (Cell Signaling), CBP80 E-7 (Santa Cruz Biotech), iEF4E FL-217 (Santa Cruz Biotech), p-4E-BP1/2/3 (Thr 45)-R (Santa Cruz Biotech) and U2AF65 MC3 (Santa Cruz Biotech).

### Cap binding experiments

Cells were stimulated with rapamycin for 24h and extracts prepared in Lysis Buffer as described above. Cell extracts were prepared in Lysis Buffer as described before. Oligo dT cellulose (Ambion) was resuspended in Lysis Buffer twice before incubating it in the presence of cell extracts on a rotating wheel at 4C during 2h. Cellulose-bound extracts were washed 5 times. RNAse treatment was done using 1/100 RNAse A/T1 cocktail (Ambion) at 37C during 13min shaking. Extracts were washed once more and cellulose was pelleted and resuspended in lysis buffer, boiled and loaded onto bis/acrylamide gels for CBP80/eIF4E blotting.

### Luciferase assays

HEK 293 T cells were starved for 24h in the absence of glutamine and then co-transfected with a normalizer firefly luciferase vector (pGL3, Promega) and pCI-Renilla/β-globin wt (WT) or NS39 (MUT) kindly gifted by Prof. A.E. Kulozik (Boelz et al. 2006). Transfected cells were then treated with vehicle (Control), 22nM of rapamycin (Rapa) or Emetine and chemiluminiscence was measured using the Dual-Luciferase Reporter (DLR) Assay (Promega) following manufacturer’s instructions.

### RT-PCR and Real Time PCR (qPCR)

Total cytoplasmic RNA and polyribosome-bound RNA was extracted using TRIzol LS following manufacturer’s instructions. In the case of polyribosomal-bound RNA, RNA was extracted from discrete polyribosomes fractions and then pooled prior to reverse transcription with random hexamers (Applied Biosystems) following manufacturer’s instructions. For actinomycin D experiments, poly-T primers were used in reverse transcription. PCR was performed using Taq Titanium (Clontech). In the case of Total to Polyribosme comparisons of CCAR1 (Fig.4) the rate of inclusion was calculated using the following formula %Inclusion= Included/(Skipped+Included) in nmol as determined using an Agilent 2001 Bioanalyzer. Statistics were done using a Mann Whitney one tailed t-test.

For qPCR experiments, Sybr Green qPCR for -PTC and +PTC was done using primers described previously in Lareau et al. 2007. Our primers were designed using the Universal Probe Library System Software (Roche) and blasted in UCSC Genome Browser tool *In silico* PCR to NCBI36/hg18 assembly:

SRSF6_All_FOR: tgg aag cag atc cag gtc tc
SRSF6_All_REV: gcg act ttt tga gat act tcg ag
CCAR1_All_FOR: CAA AAA GAA GAA CAG AAG GAG TTA GAG
CCAR1_All_REV: TCG TCT TCA GAT TTC CTA TCA TCA
CCAR1 -PTC FOR: AAC CCT CTC CCG AGG ATA CA
CCAR1_-PTC_REV: TCA TCC TTC TCT TCT TCA TCC TG
CCAR1_+PTC_FOR: TGG ACC AGA CCC AGA AAA AG
CCAR1_+PTC_REV: TTC TTC ATC CAT TGT GTG C.

Levels of each of each isoform (“-PTC” or “+PTC”) were normalized to total transcript levels (“All”) which accounts for the total gene expression of each gene and analyzed using the DDCt method. Total levels of transcript were normalized against SDHA. Statistics were done using a 2 tailed t-test and calculated as fold rapamycin-treated over control.

## RESULTS

### Depletion of PTC-containing mRNA isoforms by rapamycin is dose dependent

To assess the effects of rapamycin on NMD substrates we assayed the abundance of NMD sensitive and insensitive mRNA isoforms over a broad rapamycin concentration range by RT-qPCR. As expected, phosphorylation of 4EBP decreases with increasing rapamycin concentrations, suggesting that the activity of the mTORC1 pathway is being modulated in a dose-dependent manner (Figure 1A). We then implemented a previously described RT-qPCR assay (25) and analyzed two different NMD substrates with different modalities of AS-NMD: a) inclusion of exon 3 in the *SRSF6* mRNA which introduces a PTC and induces NMD and b) skipping of exon 13 from *CCAR1* which leads to a frameshift that triggers NMD (2). Figures 1B and 1C show the results of RT-qPCR analysis of PTC-containing and PTC-non-containing (non-PTC) isoforms for *SRSF6* and *CCAR1* genes, respectively. PTC-containing isoforms of *SRSF6* and *CCAR1* genes diminish with increasing concentrations of rapamycin. This decrease is statistically significant at the highest doses, 22nM and 2.2nM of rapamycin. We also assayed the levels of total gene expression for *SRSF6* and *CCAR1* employing primers to amplify constitutive exons in each gene and using *SDHA* as housekeeper normalizer according to (25). Our data for *SRSF6* and *CCAR1* suggest that rapamycin causes an overall increase in the total mRNA expression of both genes (Supplemental Figure 1).

**Figure 1.**
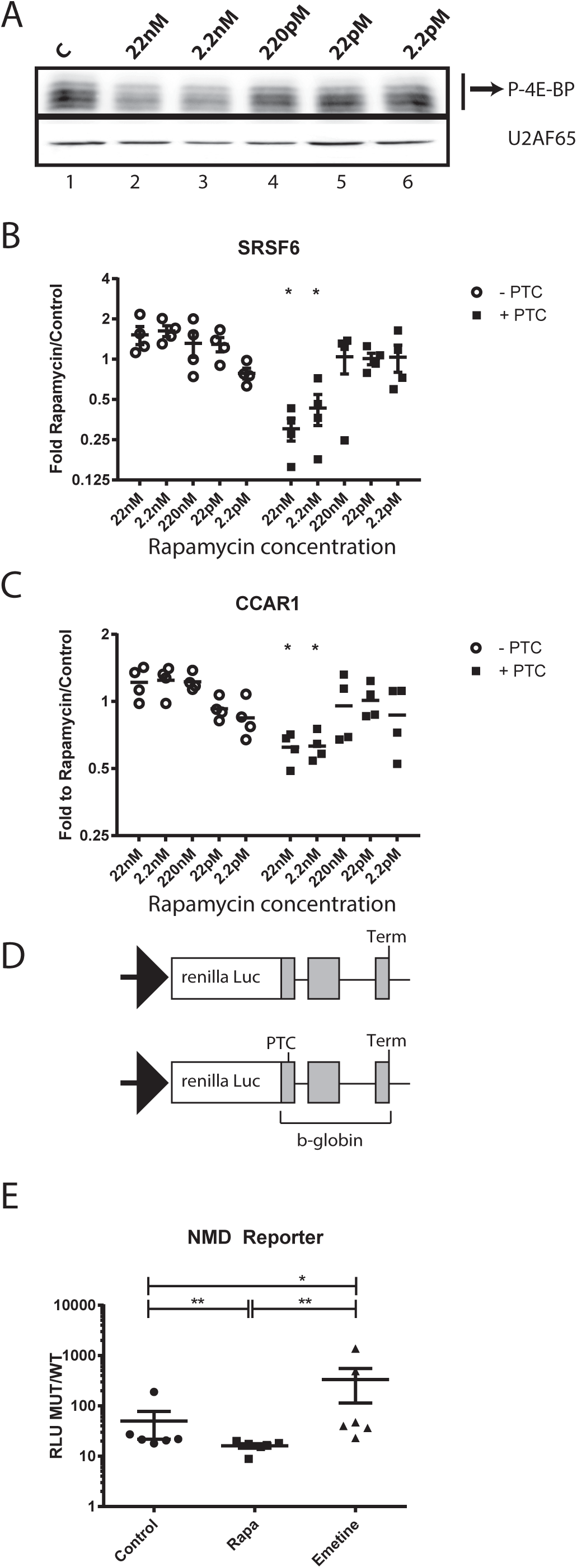
PTC-containing mRNA isoforms respond to rapamycin in a dose-dependent manner. A. Western blot analysis of 4EBP and U2AF65 (upper and lower panel) from whole cell lysates treated with decreasing concentrations of rapamycin. B. Relative RT-qPCR analysis of SRSF6 mRNA isoform (−/+ PTC) levels (normalized to total SRSF6 levels, paired 2-tailed t-test n=5) from cytoplasmic extracts of HEK 293 cells treated with rapamycin as described above. C. RT-qPCR analysis of CCAR1 mRNA isoforms (−/+, normalized to total CCAR1 mRNA levels, paired 2-tailed t-test n=5) as described above. D. Schematic for the luciferase reporter (Boeltz et al 2006) employed in 1E. E. Luciferase assays for rapamycin effects on an NMD-reporter. Y axis represents the ratio of Relative Luminescence Units of the normalized renilla NMD-reporter compared to the normalized WT-reporter (RLU MUT/WT). X axis represents the different treatments: vehicle (Control), 22nM rapamycin (Rapa) or emetine (Emetine) (n=3, Mann Whitney 2-tailed t-test).

To determine the effects of rapamycin in exogenous PTC-containing transcripts we employed a previously developed and validated NMD reporter (26). We co-transfected HEK 293 T cells with a reporter vector containing a wild type -globin open reading frame (WT) or an NMD-triggering mutation -globin (MUT) fused to a renilla luciferase gene (schematics in Figure 1D) together with a normalizer firefly luciferase vector. Transfected cells were then treated with vehicle (Control), 22nM of rapamycin (Rapa) or emetine (Emetine) and chemiluminiscence was measured. We employed emetine as a control, because it inhibits NMD leading to up-regulation of PTC-containing transcripts (27). Figure 1E shows our results as Relative Luminescence Units of the normalized renilla NMD-reporter compared to the normalized WT-reporter (RLU MUT/WT) as in (26). Our data indicate that the expression of the NMD reporter was significantly diminished in cells treated with rapamycin compared to vehicle (compare Rapa to Control). As expected, emetine treatment increased the stability of the NMD reporter. Taken together, our data suggest a widespread mechanism for rapamycin triggering depletion of PTC-containing mRNAs.

### Rapamycin affects mRNA stability of PTC-containing mRNA isoforms and requires UPF1

The previous experiments suggested that rapamycin augments NMD activity and decreases levels of PTC-containing isoforms in HEK cells. In order to confirm that the observed change in the ratio of PTC-containing/non-PTC isoforms is due to RNA decay and not merely AS regulation, we assayed the half-life of PTC-containing and non-PTC isoforms of *SRSF6* in a rapamycin dependent manner using RT-qPCR. Figure 2A shows that endogenous *SRSF6* PTC-containing isoform displayed accelerated decay kinetics in rapamycin-treated cells as compared to the non-PTC isoform, suggestive of an increased rate of NMD rather than regulation of AS.

**Figure 2.**
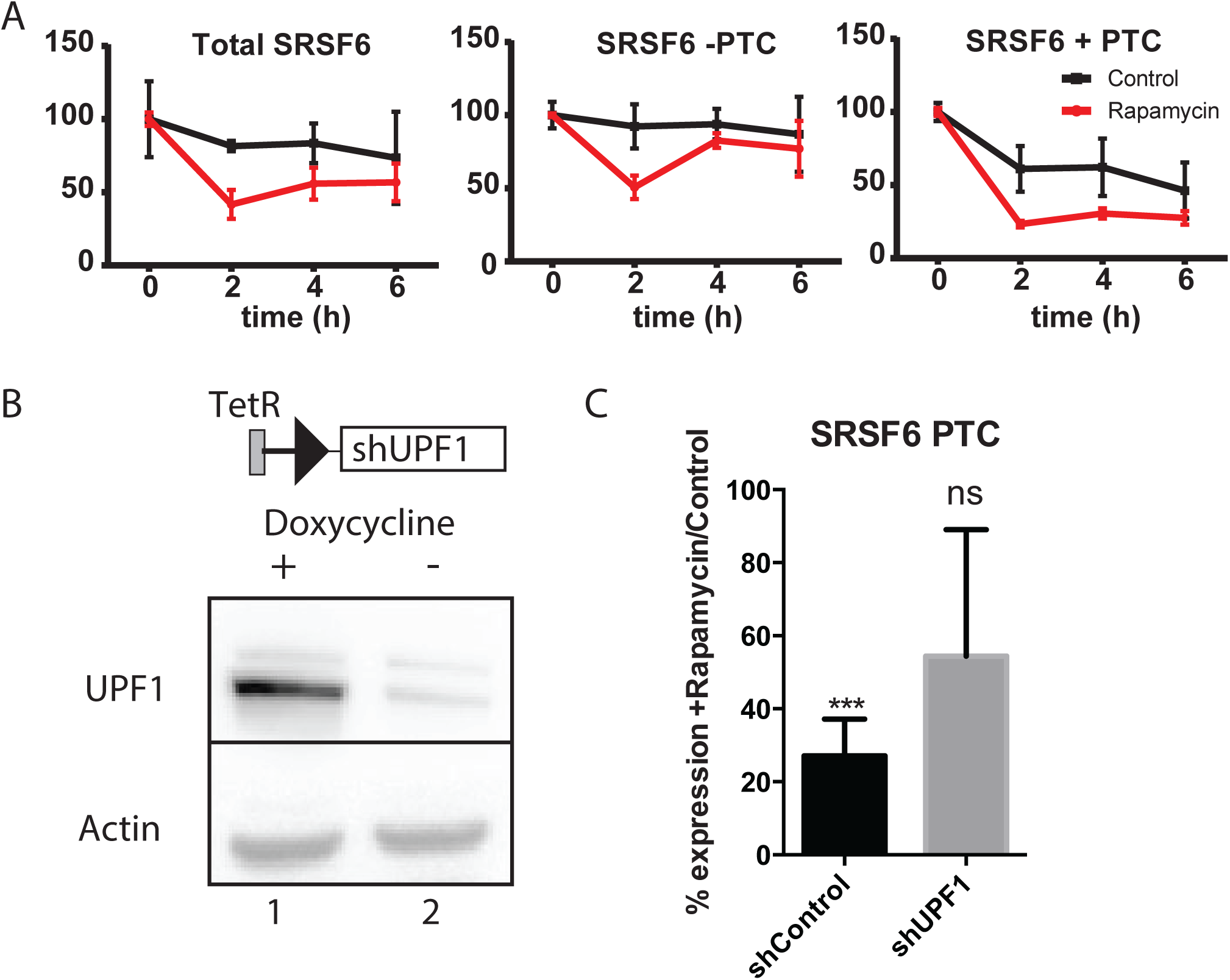
Rapamycin alters the stability of PTC-containing isoforms and its effects are UPF1-dependent. A. Relative RT-qPCR analysis of constitutive SRSF6, SRSF6 non-PTC isoforms (SRSF6 -PTC) and PTC-containing (SRSF6 +PTC) after rapamycin treatment and translational arrest with actinomycin D to measure mRNA half life (n=5). B. Representative western blot of the inducible Tet-on shUPF1 cell line. C. RT-qPCR analysis of SRSF6 PTC+ mRNA isoforms in control or UPF1-depleted cells treated with or without rapamycin (n=4).

To test the direct role of NMD machinery in rapamycin effects, we compared the levels of PTC-containing *SRSF6* mRNA isoforms in UPF1-depleted HEK cells treated with or without rapamycin. Depletion of UPF1 inhibits NMD in a variety of model systems (28,29). We generated stably transduced HEK cell lines with an inducible Tet-On shRNA against UPF1 or a scramble control (30) (Figure 2B). As previously observed, rapamycin stimulation decreased the levels of PTC-containing isoforms of endogenous SRSF6 in control cells. However, this effect was attenuated in the UPF1 depleted cells (Figure 2C). These data demonstrate that the depletion of PTC-containing isoforms by rapamycin is dependent on UPF1 and supports our hypothesis that rapamycin enhances NMD.

### Rapamycin decreases levels of PTC-containing isoforms bound to polyribosomes

NMD can happen not only during pioneer translation but also during steady translation, given that eIF4E-bound mRNAs are substrates of NMD (19). Moreover, Durand *et al.* demonstrated that eIF4F-bound transcripts are susceptible to NMD, even when translation is inhibited (18). A prediction from these experiments is that NMD efficiency of polyribosome-associated mRNAs may be dynamically regulated. Given that NMD in mammals is translation-dependent (31,32) and that rapamycin is a global translation modulator, we hypothesized that rapamycin-treatment might enhance the decay of polyribosome-bound PTC-containing transcripts. We investigated this hypothesis in cells treated with or without rapamycin. As expected, rapamycin-treatment increased the electrophoretic mobility of 4EBP on SDS-PAGE relative to control cells (Figure 3A), indicative of 4EBP hypophosphorylation. We then isolated polyribosomes from control or rapamycin treated cells. We observed that rapamycin treated cells showed the distinctive characteristics of global translational inhibition when compared to control cells: an increase in the monosomal fraction (80S, peak) and a decrease of the polyribosome fractions (Figure 3B shows a representative polyribosome profile). We then analyzed the presence of PTC-containing and non-PTC isoforms in the polyribosomal fractions. In control cells we observed an enrichment of the *SRSF6* PTC-containing isoform in the polyribosome-associated mRNA fraction as compared to total cytoplasmic mRNAs (Figure 3C, compare lanes 1 and 2). Rapamycin-treatment dramatically reduced levels of the PTC-containing *SRSF6* isoform in polyribosome-associated mRNA fractions (Figure 3C, compare lanes 2 and 4). To rule out alternative splicing as the underlying cause of PTC-containing isoform depletion from polyribosomes in rapamycin-treated cells, we stimulated cells with both rapamycin and the translation elongation inhibitor emetine. We expected that if rapamycin effects were splicing dependent, then emetine treatment would not alter the levels of PTC-containing isoforms observed in polyribosomal-bound fractions between control and rapamycin treated cells. As expected, we observed that the effect of rapamycin was attenuated by emetine (compare lanes 6 and 8 in Figure 3C). Taken together these data suggest that rapamycin alters the stability of PTC-containing isoforms in polyribosome complexes rather than the AS of their pre-mRNA.

**Figure 3.**
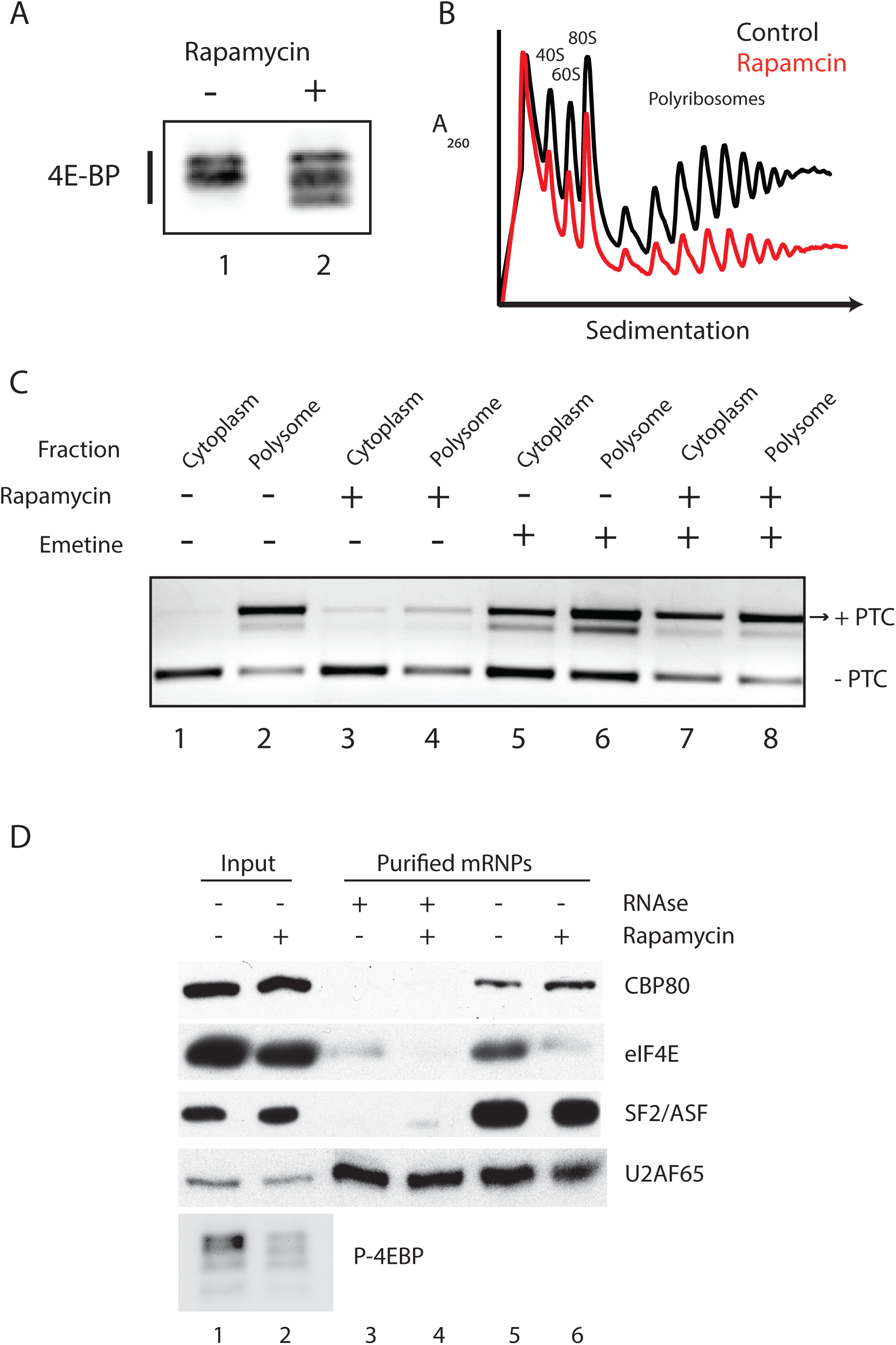
Rapamycin alters the levels of polyribosome associated PTC-containing mRNA isoforms. A. Representative western blot of 4EBP levels from control or rapamycin-treated cells. B. Representative 260 nm UV absorbance across 15-45% sucrose gradients of cytoplasmic extracts from cells treated with or without rapamycin (n=3). C. Endpoint RT-PCR analysis of PTC+/− SRSFF6 mRNA isoforms in either cytoplasmic (T) or polyribosome associated (P) extracts treated with or without rapamycin and emetine. D. Western blot analysis of proteins co-purified with poly A+ RNA from cytoplasmic extracts of control or rapamycin-treated cells.

We further assayed the abundance of PTC-containing and non-PTC isoforms of *SRSF6* and *CCAR1* in total and polyribosome-bound mRNA fractions at 7h post-rapamycin stimulation. Rapamycin significantly diminished the presence of the PTC-containing isoforms in polyribosomal fractions of both SFRS6 and CCAR1 (Supplemental Figure 2) genes whereas the effects were not detectable when comparing the total pool of mRNAs. Together these data suggest that rapamycin diminishes the abundance of PTC-containing transcripts through a translation-dependent mechanism.

Previous studies demonstrated that rapamycin treatment induces nuclear accumulation of eIF4E through de-phosphorylation and de-repression of 4EBP (16,33). We confirmed these data in HEK 293 T cells: upon glutamine starvation we observed rapamycin-dependent modest nuclear accumulation of eIF4E whereas it is localized predominantly to the cytoplasm in control cells. In contrast, in rapamycin-treated cells we observed strong accumulation of eIF4E in the nucleus (Supplemental Figure 3). Given our previous results suggesting a role for rapamycin in the degradation of PTC-containing isoforms, and because rapamycin treatment causes nuclear sequestration of eIF4E, we hypothesized that exchange of cap binding proteins on mRNAs may also be abrogated. We thus purified cytoplasmic mRNPs from control or rapamycin-treated cells and assayed eIF4E and CBP80 association in an RNA-dependent manner. Figure 3D shows a representative western blot of eIF4E, CBP80, SF2/ASF, and U2AF65 in whole cell extracts (lanes 1 and 2) and RNase-treated or untreated eluates from oligo dT cellulose affinity chromatography. We observed that rapamycin-treatment increases the electrophoretic mobility of 4EBP, consistent with de-phosphorylation. With the exception of U2AF65, which binds directly to oligo dT cellulose, SF2/ASF, CBP80 and eIF4E were all RNase-sensitive, suggesting that their binding to oligo dT cellulose is RNA-dependent. Unlike SF2/ASF protein, which bound to polyA-mRNAs equally in the presence or absence of rapamycin, CBP80 and eIF4E mRNPs levels were inversely correlated (Figure 3D lanes 5-6). Treatment with rapamycin resulted in a decrease of eIF4E association with polyadenylated mRNPs and a modest but reproducible increase in CBP80 association. Together, our data suggests that rapamycin-treatment impairs the normal exchange of CBP80 with eIF4E on cytoplasmic mRNPs, rendering them more susceptible to nonsense mediated decay and subsequent increased degradation of PTC-containing mRNAs.

## DISCUSSION

Little is known about enhancement of NMD, which may have the potential of increasing genomic surveillance by increasing the degradation of potentially deleterious mRNAs. Up to 11% of genetic inherited diseases are due to deleterious NMD (34) underlining the importance of NMD. The accumulation of potentially “toxic” mRNAs can lead to translation of truncated proteins that may be harmful for the cells, especially under stress conditions. As an example, the only hallmark of pancreatic adenosquamous carcinoma is a deficient NMD signature due to mutations in UPF1 (35). These data suggest that genomic surveillance is required for a “healthy” cellular environment and the possibility that cells have evolved a mechanism to integrate danger signals coming from mRNAs. It is also plausible to consider the idea that accumulation of “toxic” mRNAs may increase cellular stress (36).

Our data demonstrate a role for rapamycin in augmenting decay of PTC-containing mRNA isoforms. Rapamycin increases the nuclear accumulation of eIF4E (Supplemental Figure 3), which is consistent with arrest of translation initiation (Figure 3B). With eIF4E being predominantly localized to the nucleus upon rapamycin exposure, mRNAs may be pushed towards a pioneer-like mRNP complex in the cytoplasm. This hypothesis is reflected in Figure 3D which strongly suggests that rapamycin impairs the exchange of the CBC component CBP80 for eIF4E favoring an increased ratio of CBC/eIF4E binding. This can in turn lead to an increased opportunity for PTC-containing mRNAs to be flagged for degradation by NMD. It is possible that in addition to this “pro-pioneer round” effect exerted by rapamycin, PTC-containing transcripts bound to eIF4E decrease in abundance due to-the nuclear accumulation of eIF4E and -the decrease of eIF4E binding to polyadenylated mRNAs upon rapamycin stimulation (model in Figure 4). Importantly, both Rufener and Durand studies pose a central role for translation in NMD (18,19), strongly supporting our observations that PTC-containing isoforms are less abundant in polyribosomes in rapamycin treated cells (Figures 3C and Supplemental Figure 2) and consistent with nonsense-mediated mRNA decay occurring during translation (31,32).

**Figure 4.**
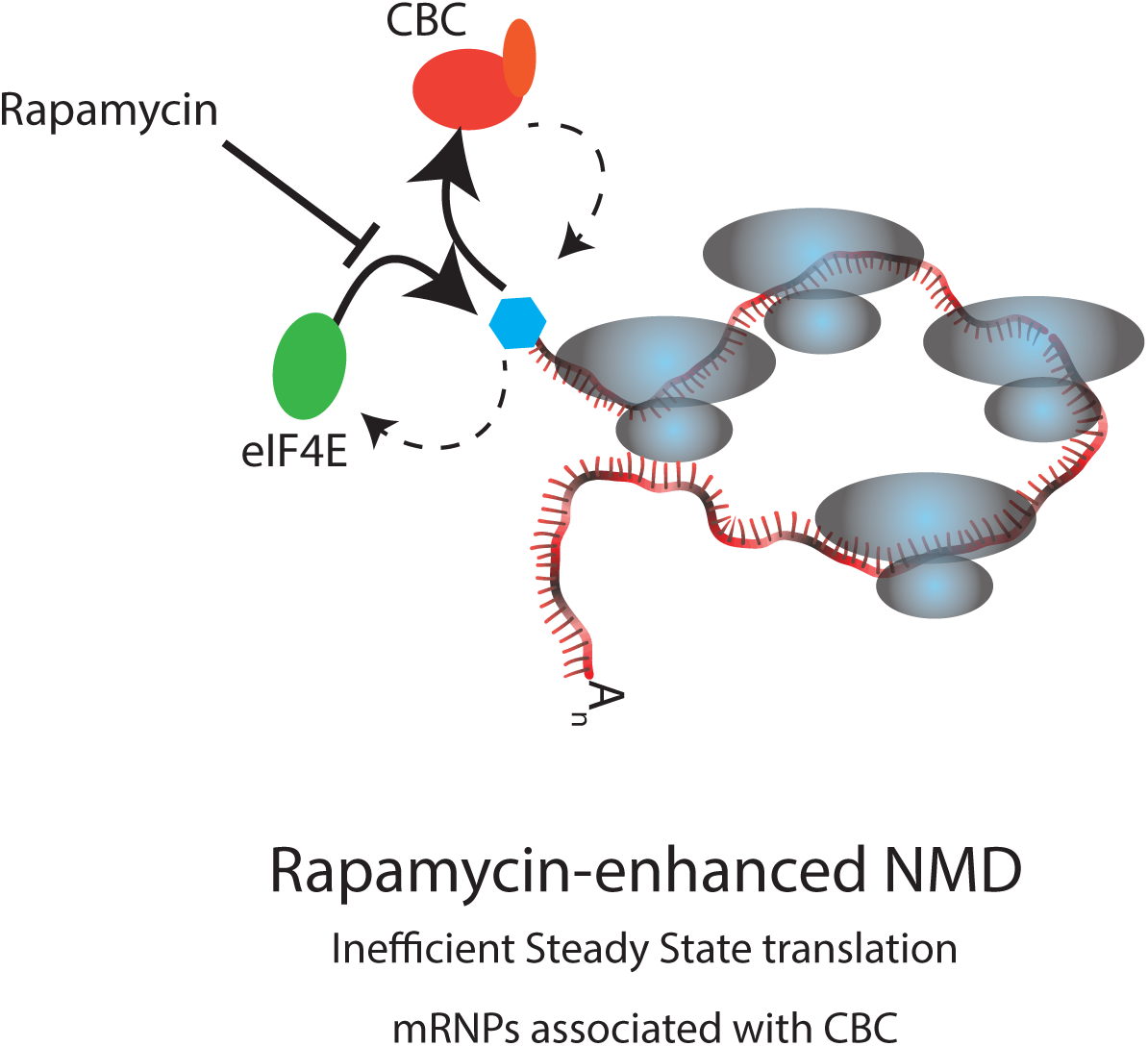
Our proposed model for rapamycin effects on NMD.

Rapamycin’s effects on NMD may also rely on post-translational modifications (such as phosphorylation) of NMD factors in addition to augmentation of CBC-binding to mRNPs (Figure 3D). Work by Pal *et al.* has shown that phosphorylation of UPF1 is sensitive to rapamycin (37); however the effect of lowered UPF1 phosphorylation on PTC-containing mRNAs was not tested. Moreover, although rapamycin concentration was similar to our working dose (25nM and 22nM, respectively), cells were only exposed for a short period of time (30min *vs* 72h in our case). We do not exclude the possibility that prolonged exposure to rapamycin may result in phosphorylation changes in some of NMD factors (or their regulators).

Augmentation of NMD by rapamycin could have broad implications for cell biology and physiology. Given that rapamycin increases lifespan in a wide variety of model systems, including mammals (38,39), it is tempting to hypothesize that by increasing NMD efficiency, rapamycin may provide a ‘healthier’ transcriptome via degradation of potentially deleterious mRNAs that would otherwise result in cellular toxicity and premature senescence. NMD is effective not only during pioneer-but also during steady-state translation. Cells may then stand a better chance for surveillance and regulation of their transcriptome. For example, a previous study showed that UPF1 levels play a key role in deciding between undifferentiated cells (high UPF1 expression) *vs* neurogenic cells (UPF1 down-regulated) (40). These data may also implicate that NMD efficiency may allow neuronal regeneration, which is depleted in age-related diseases. We believe our study presents novel experimental evidence for the role of rapamycin in NMD that may have a broad impact in ageing and age-related diseases.

## ACKNOWLEDGEMENT

The authors thank Javier Caceres, Luiz Penalva, Alan Zahler and Aishwarya Griselda Jacob for critical input to the manuscript. The authors also thank Prof. A.E. Kulozik for kindly providing pCI-Renilla/β-globin wt (WT) or NS39 (MUT) to perform the luciferase experiments.

---

**Supplemental Figure 1.**
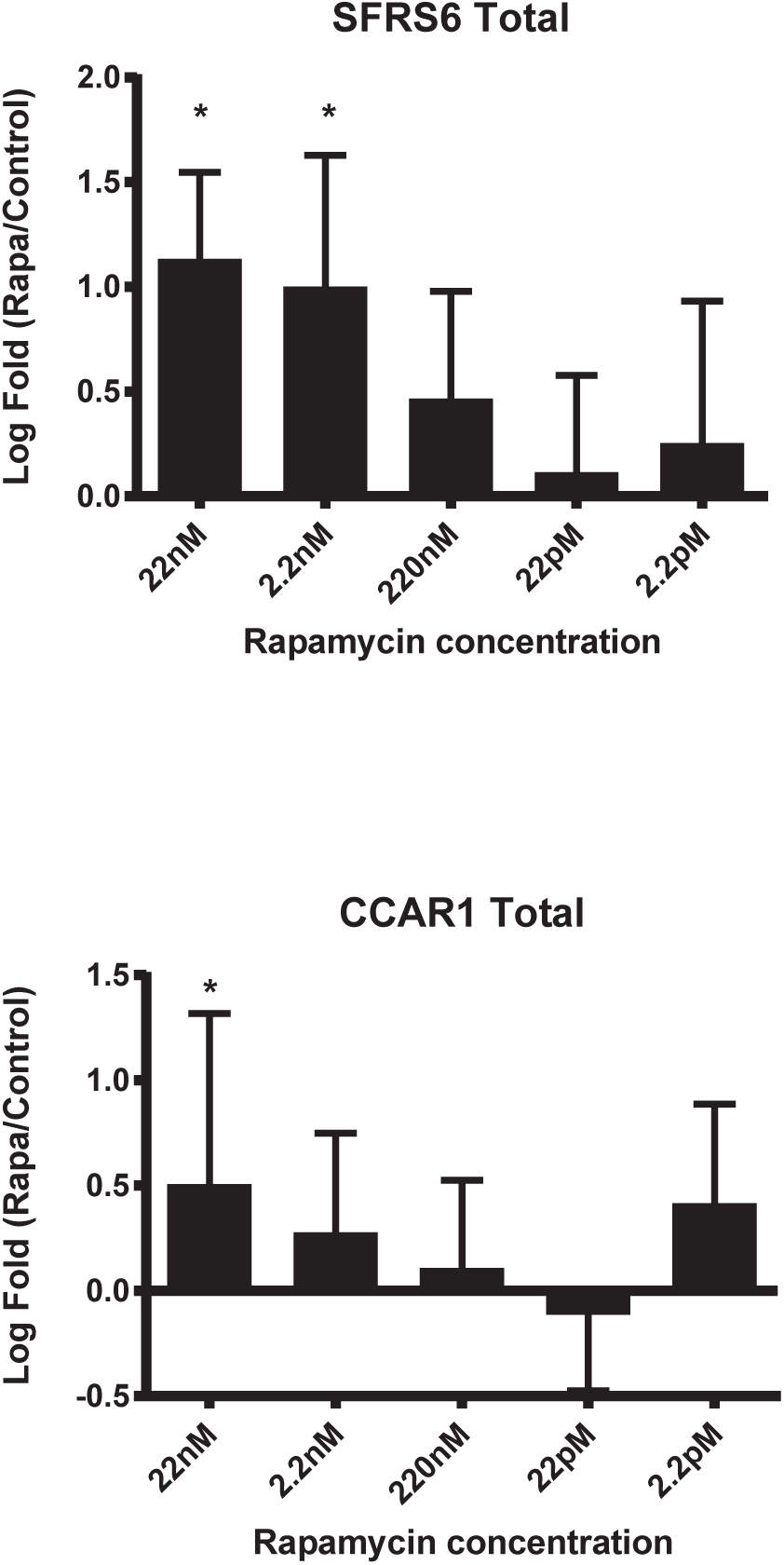
A. Relative RT-qPCR analysis of SRSF6 mRNA levels (normalized to SDHA levels, n=5) from cytoplasmic extracts of HEK 293 cells treated with rapamycin as described above. B. RT-qPCR analysis of CCAR1 mRNA normalized to total SDHA mRNA levels, n=5) as described above.

**Supplemental Figure 2.**
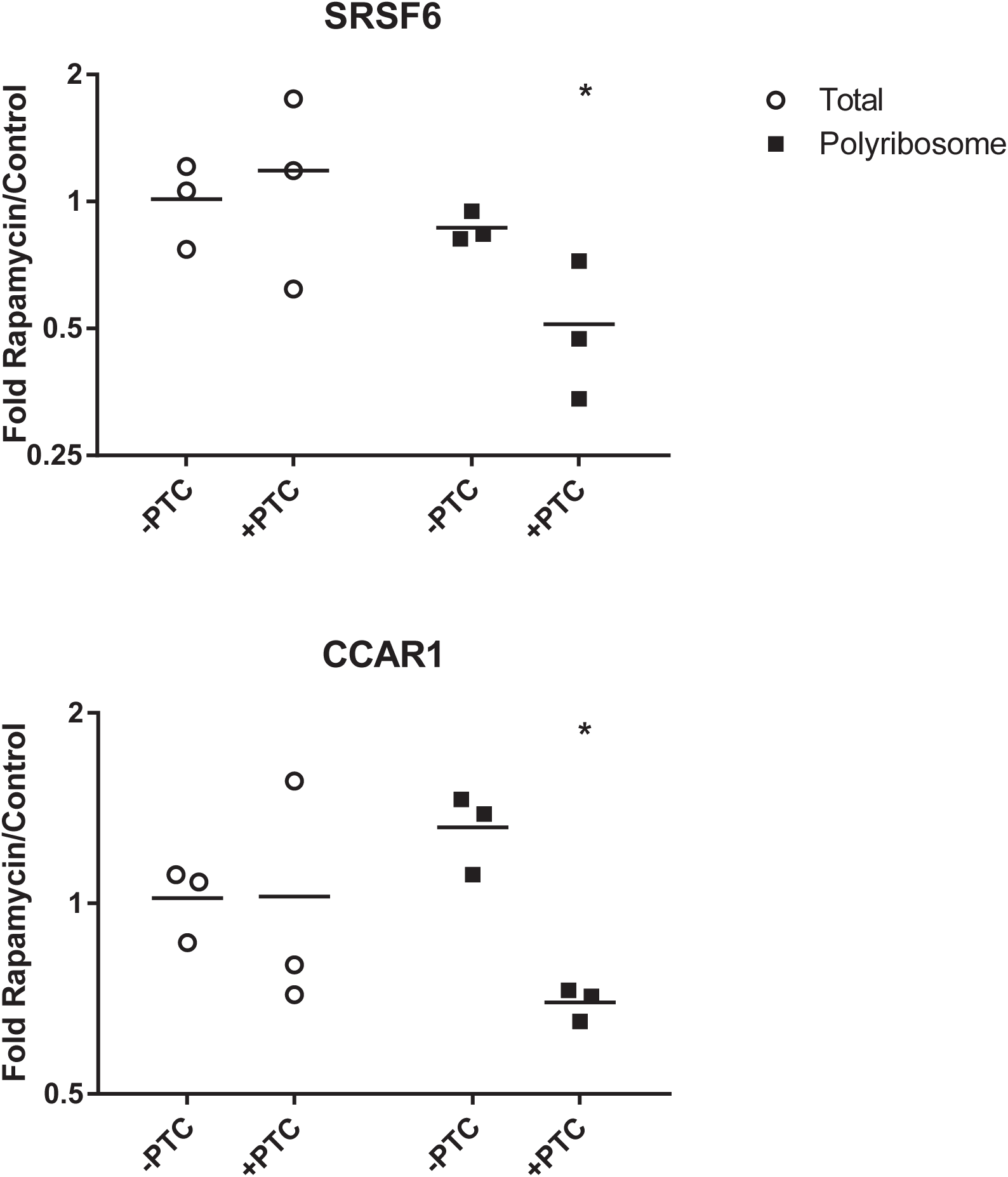
Rapamycin alters the subcellular localization of eIF4E in HEK293 cells. A. Representative immunoflourescence micrograph of HEK 293 cells stained with antibodies against eIF4E and counter stained with DAPI. Cells were starved (no glutamine), Fed after starvation (glutamine) or fed in the presence of rapamycin (Gln, Rapamycin). B. Quantification of eIF4E staining across individual cells.

**Supplemental Figure 3.**
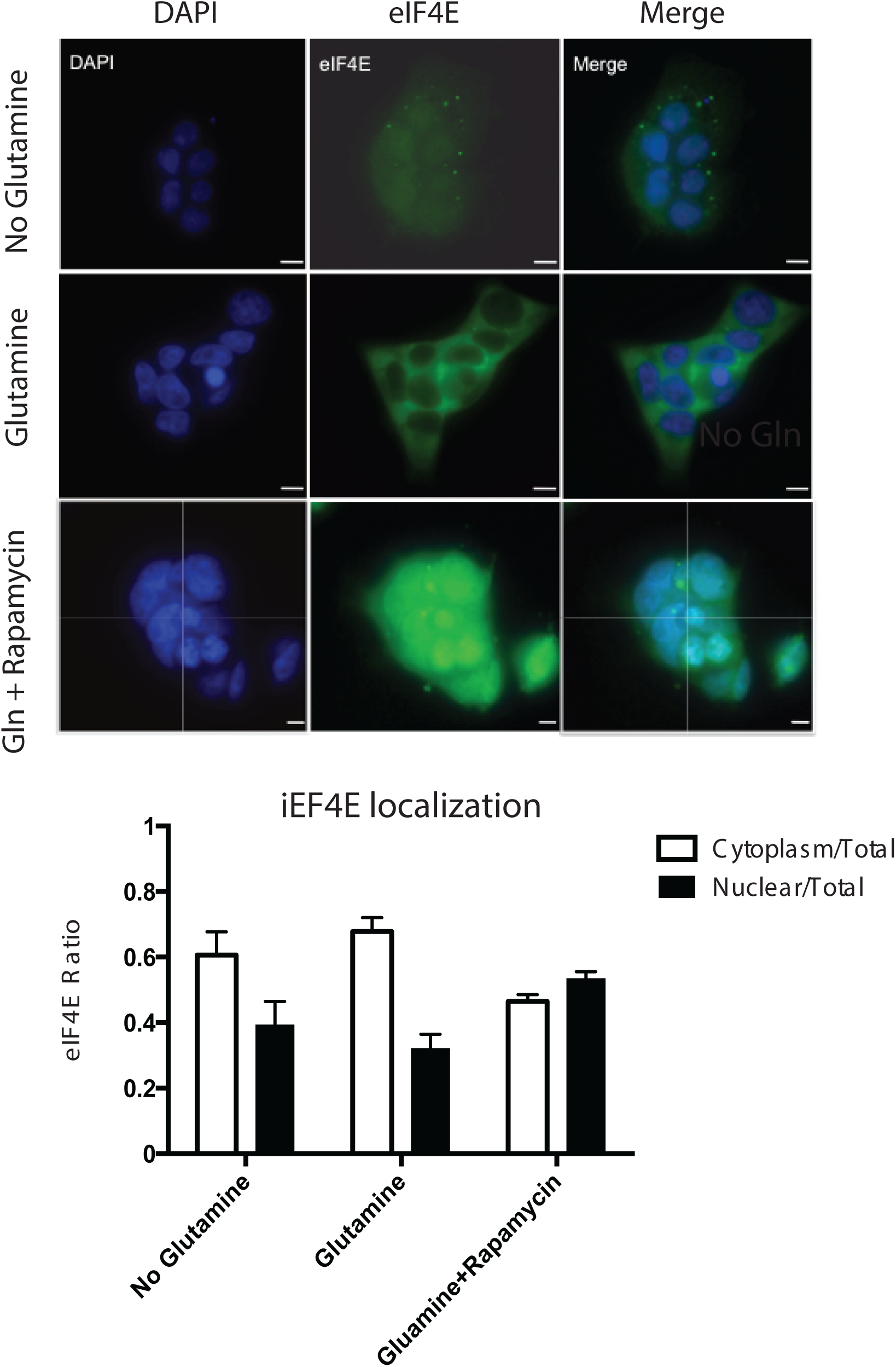
RT-qPCR analysis of SRSF6 and CCAR1 PTC-containing and non-containing mRNA isoforms (normalized to total respective mRNA levels) in cytoplasmic or polyribosomal extracts prepared from cells treated with or without rapamycin (P=0.0143 and P=0.05, respectively, Mann-Whitney test, n=3).

## REFERENCES

1. Popp, M.W. and Maquat, L.E. (2013) Organizing principles of mammalian nonsense-mediated mRNA decay. Annual review of genetics, 47, 139–165.

2. Ni, J.Z., Grate, L., Donohue, J.P., Preston, C., Nobida, N., O’Brien, G., Shiue, L., Clark, T.A., Blume, J.E. and Ares, M., Jr. (2007) Ultraconserved elements are associated with homeostatic control of splicing regulators by alternative splicing and nonsense-mediated decay. Genes & development, 21, 708–718.

3. Huang, L., Lou, C.H., Chan, W., Shum, E.Y., Shao, A., Stone, E., Karam, R., Song, H.W. and Wilkinson, M.F. (2011) RNA homeostasis governed by cell type-specific and branched feedback loops acting on NMD. Molecular cell, 43, 950–961.

4. Le Hir, H., Izaurralde, E., Maquat, L.E. and Moore, M.J. (2000) The spliceosome deposits multiple proteins 20-24 nucleotides upstream of mRNA exon-exon junctions. The EMBO journal, 19, 6860–6869.

5. Ishigaki, Y., Li, X., Serin, G. and Maquat, L.E. (2001) Evidence for a pioneer round of mRNA translation: mRNAs subject to nonsense-mediated decay in mammalian cells are bound by CBP80 and CBP20. Cell, 106, 607–617.

6. Lejeune, F., Ishigaki, Y., Li, X. and Maquat, L.E. (2002) The exon junction complex is detected on CBP80-bound but not eIF4E-bound mRNA in mammalian cells: dynamics of mRNP remodeling. The EMBO journal, 21, 3536–3545.

7. Dostie, J. and Dreyfuss, G. (2002) Translation is required to remove Y14 from mRNAs in the cytoplasm. Current biology: CB, 12, 1060–1067.

8. Sato, H. and Maquat, L.E. (2009) Remodeling of the pioneer translation initiation complex involves translation and the karyopherin importin beta. Genes & development, 23, 2537–2550.

9. Lejeune, F., Li, X. and Maquat, L.E. (2003) Nonsense-mediated mRNA decay in mammalian cells involves decapping, deadenylating, and exonucleolytic activities. Molecular cell, 12, 675–687.

10. Chiu, S.Y., Lejeune, F., Ranganathan, A.C. and Maquat, L.E. (2004) The pioneer translation initiation complex is functionally distinct from but structurally overlaps with the steady-state translation initiation complex. Genes & development, 18, 745–754.

11. Kashima, I., Yamashita, A., Izumi, N., Kataoka, N., Morishita, R., Hoshino, S., Ohno, M., Dreyfuss, G. and Ohno, S. (2006) Binding of a novel SMG-1-Upf1-eRF1-eRF3 complex (SURF) to the exon junction complex triggers Upf1 phosphorylation and nonsense-mediated mRNA decay. Genes & development, 20, 355–367.

12. Ohnishi, T., Yamashita, A., Kashima, I., Schell, T., Anders, K.R., Grimson, A., Hachiya, T., Hentze, M.W., Anderson, P. and Ohno, S. (2003) Phosphorylation of hUPF1 induces formation of mRNA surveillance complexes containing hSMG-5 and hSMG-7. Molecular cell, 12, 1187–1200.

13. Eberle, A.B., Stalder, L., Mathys, H., Orozco, R.Z. and Muhlemann, O. (2008) Posttranscriptional gene regulation by spatial rearrangement of the 3’ untranslated region. PLoS biology, 6, e92.

14. Toma, K.G., Rebbapragada, I., Durand, S. and Lykke-Andersen, J. (2015) Identification of elements in human long 3’ UTRs that inhibit nonsense-mediated decay. Rna.

15. Dias, S.M., Wilson, K.F., Rojas, K.S., Ambrosio, A.L. and Cerione, R.A. (2009) The molecular basis for the regulation of the cap-binding complex by the importins. Nature structural & molecular biology, 16, 930–937.

16. Rong, L., Livingstone, M., Sukarieh, R., Petroulakis, E., Gingras, A.C., Crosby, K., Smith, B., Polakiewicz, R.D., Pelletier, J., Ferraiuolo, M.A. et al. (2008) Control of eIF4E cellular localization by eIF4E-binding proteins, 4E-BPs. Rna, 14, 1318–1327.

17. Hosoda, N., Kim, Y.K., Lejeune, F. and Maquat, L.E. (2005) CBP80 promotes interaction of Upf1 with Upf2 during nonsense-mediated mRNA decay in mammalian cells. Nature structural & molecular biology, 12, 893–901.

18. Durand, S. and Lykke-Andersen, J. (2013) Nonsense-mediated mRNA decay occurs during eIF4F-dependent translation in human cells. Nature structural & molecular biology, 20, 702–709.

19. Rufener, S.C. and Muhlemann, O. (2013) eIF4E-bound mRNPs are substrates for nonsense-mediated mRNA decay in mammalian cells. Nature structural & molecular biology, 20, 710–717.

20. Kim, S.G., Buel, G.R. and Blenis, J. (2013) Nutrient regulation of the mTOR complex 1 signaling pathway. Molecules and cells, 35, 463–473.

21. Laplante, M. and Sabatini, D.M. (2012) mTOR signaling in growth control and disease. Cell, 149, 274–293.

22. Gingras, A.C., Raught, B. and Sonenberg, N. (2001) Regulation of translation initiation by FRAP/mTOR. Genes & development, 15, 807–826.

23. Hsieh, A.C., Liu, Y., Edlind, M.P., Ingolia, N.T., Janes, M.R., Sher, A., Shi, E.Y., Stumpf, C.R., Christensen, C., Bonham, M.J. et al. (2012) The translational landscape of mTOR signalling steers cancer initiation and metastasis. Nature, 485, 55–61.

24. Joncourt, R., Eberle, A.B., Rufener, S.C. and Muhlemann, O. (2014) Eukaryotic initiation factor 4G suppresses nonsense-mediated mRNA decay by two genetically separable mechanisms. PloS one, 9, e104391.

25. Lareau, L.F., Inada, M., Green, R.E., Wengrod, J.C. and Brenner, S.E. (2007) Unproductive splicing of SR genes associated with highly conserved and ultraconserved DNA elements. Nature, 446, 926–929.

26. Boelz, S., Neu-Yilik, G., Gehring, N.H., Hentze, M.W. and Kulozik, A.E. (2006) A chemiluminescence-based reporter system to monitor nonsense-mediated mRNA decay. Biochemical and biophysical research communications, 349, 186–191.

27. Noensie, E.N. and Dietz, H.C. (2001) A strategy for disease gene identification through nonsense-mediated mRNA decay inhibition. Nature biotechnology, 19, 434–439.

28. Hurt, J.A., Robertson, A.D. and Burge, C.B. (2013) Global analyses of UPF1 binding and function reveal expanded scope of nonsense-mediated mRNA decay. Genome research, 23, 1636–1650.

29. Leeds, P., Peltz, S.W., Jacobson, A. and Culbertson, M.R. (1991) The product of the yeast UPF1 gene is required for rapid turnover of mRNAs containing a premature translational termination codon. Genes & development, 5, 2303–2314.

30. Bulliard, Y., Wiznerowicz, M., Barde, I. and Trono, D. (2006) KRAB can repress lentivirus proviral transcription independently of integration site. The Journal of biological chemistry, 281, 35742–35746.

31. Hu, W., Petzold, C., Coller, J. and Baker, K.E. (2010) Nonsense-mediated mRNA decapping occurs on polyribosomes in Saccharomyces cerevisiae. Nature structural & molecular biology, 17, 244–247.

32. Hu, W., Sweet, T.J., Chamnongpol, S., Baker, K.E. and Coller, J. (2009) Co-translational mRNA decay in Saccharomyces cerevisiae. Nature, 461, 225–229.

33. Dostie, J., Ferraiuolo, M., Pause, A., Adam, S.A. and Sonenberg, N. (2000) A novel shuttling protein, 4E-T, mediates the nuclear import of the mRNA 5’ cap-binding protein, eIF4E. The EMBO journal, 19, 3142–3156.

34. Mort, M., Ivanov, D., Cooper, D.N. and Chuzhanova, N.A. (2008) A meta-analysis of nonsense mutations causing human genetic disease. Human mutation, 29, 1037–1047.

35. Liu, C., Karam, R., Zhou, Y., Su, F., Ji, Y., Li, G., Xu, G., Lu, L., Wang, C., Song, M. et al. (2014) The UPF1 RNA surveillance gene is commonly mutated in pancreatic adenosquamous carcinoma. Nature medicine, 20, 596–598.

36. Gardner, L.B. (2010) Nonsense-mediated RNA decay regulation by cellular stress: implications for tumorigenesis. Molecular cancer research: MCR, 8, 295–308.

37. Pal, M., Ishigaki, Y., Nagy, E. and Maquat, L.E. (2001) Evidence that phosphorylation of human Upfl protein varies with intracellular location and is mediated by a wortmannin-sensitive and rapamycin-sensitive PI 3-kinase-related kinase signaling pathway. Rna, 7, 5–15.

38. Bjedov, I., Toivonen, J.M., Kerr, F., Slack, C., Jacobson, J., Foley, A. and Partridge, L. (2010) Mechanisms of life span extension by rapamycin in the fruit fly Drosophila melanogaster. Cell metabolism, 11, 35–46.

39. Harrison, D.E., Strong, R., Sharp, Z.D., Nelson, J.F., Astle, C.M., Flurkey, K., Nadon, N.L., Wilkinson, J.E., Frenkel, K., Carter, C.S. et al. (2009) Rapamycin fed late in life extends lifespan in genetically heterogeneous mice. Nature, 460, 392–395.

40. Lou, C.H., Shao, A., Shum, E.Y., Espinoza, J.L., Huang, L., Karam, R. and Wilkinson, M.F. (2014) Posttranscriptional control of the stem cell and neurogenic programs by the nonsense-mediated RNA decay pathway. Cell reports, 6, 748–764.

